# Early induction of hepatic deiodinase type 1 inhibits hepatosteatosis during NAFLD progression

**DOI:** 10.1101/2021.05.13.443791

**Authors:** Eveline Bruinstroop, Jin Zhou, Madhulika Tripathi, Winifred W. Yau, Anita Boelen, Brijesh Kumar Singh, Paul M. Yen

## Abstract

**Objective:** Non-alcoholic fatty liver disease (NAFLD) comprises a spectrum ranging from hepatosteatosis to progressive non-alcoholic steatohepatitis that can lead to cirrhosis. Humans with low levels of the prohormone thyroxine (T_4_) have a higher incidence of NAFLD and thyroid hormone treatment is very promising in all patients with NAFLD. Deiodinase 1 (Dio1) is a hepatic enzyme that converts T_4_ to the bioactive T_3_, and therefore regulates intracellular thyroid hormone availability in the liver. The role of this intracellular regulation was investigated during the progression of NAFLD.

**Methods:** We investigated intracellular thyroid hormone metabolism in two NAFLD models: wildtype mice fed Western diet with fructose and *Lepr*^db^ mice fed a methionine- and choline-deficient diet. AAV8-mediated liver-specific Dio1 knockdown was employed to investigate the role of Dio1 during the progression of NAFLD. Intrahepatic thyroid hormone levels, deiodinase activity and metabolic parameters were measured.

**Results:** Dio1 expression and activity were increased in the early stages of NAFLD and were associated with an increased T_3_/T_4_ ratio. Prevention of this increase by AAV8-mediated liver-specific Dio1 knockdown increased hepatic triglycerides and cholesterol as well as decreased pACC/ACC ratio and acylcarnitine levels suggesting there was lower β-oxidation. Dio1 siRNA KD in hepatic cells treated with fatty acids showed increased lipid accumulation and decreased oxidative phosphorylation.

**Conclusion:** Hepatic Dio1 gene expression was modulated by dietary conditions, increased during hepatosteatosis and early NASH, and regulated hepatic triglyceride content. These early adaptations likely represent compensatory mechanisms to reduce hepatosteatosis and prevent NASH progression.

## 1. Introduction

Non-alcoholic fatty liver disease (NAFLD) comprises a spectrum of disease ranging from simple steatosis in the liver to steatohepatitis (NASH) with inflammation and fibrosis. NAFLD affects approximately 25% of the adult population worldwide, and its rise has been associated with the recent pandemic of obesity and diabetes [1]. Currently, there are no approved drugs for the treatment of NAFLD so there is an urgent need for the development of new therapies. Recently, thyroid hormone (TH) and TH-analogs (thyromimetics) have shown to be effective therapies for hepatosteatosis and NASH [2–6]. However, the physiological basis of their effects on NAFLD are not well understood.

TH stimulates lipophagy, β-oxidation of fatty acids, and oxidative phosphorylation in the liver [4 7]. Previous studies showed that both hypothyroidism and thyroid hormone receptor β mutations in mouse and man increase the risk of NAFLD [8 9]. Also, lower serum levels of prohormone thyroxine (T_4_), even within the normal range increase the prevalence of NAFLD [10]. However, serum levels of T_4_ are not the only factor determining intracellular concentrations of the bioactive triiodothyronine (T_3_) which binds to the nuclear hormone receptor β (TRβ) causing transcriptional activation of T_3_ target genes. Intracellular concentrations of the prohormone T_4_ and the bioactive form of TH, T_3_, are tightly regulated by intracellular deiodinases. There are three deiodinases, deiodinase type1 (Dio1), deiodinase type 2 (Dio2) and deiodinase type 3 (Dio3), all selenoenzymes of which Dio1 and Dio3 are expressed in hepatocytes. Dio1 is responsible for outer and inner ring deiodination of thyroid hormone and thereby involved in T_3_ production and rT_3_ clearance. Dio3 regulates T_3_ and T_4_ conversion to the inert metabolites T_2_ and rT_3_ respectively. The expression of Dio1 is influenced by cytokines and nutritional status, and it is markedly up-regulated by T_3_ [2 11]. Previously, low levels of Dio1 expression were found in the livers of mice after acute and chronic inflammation, as well as in patients with NASH [12 13]. In this study, we examined the role of intracellular regulation of thyroid hormone during the different phases of NAFLD progression.

## 2. Materials and Methods

### 2.1 General

All mice were maintained according to the Guide for the Care and Use of Laboratory Animals [National Institutes of Health (NIH) publication 1.0.0; revised 2011], and experiments were approved by the Singhealth Institutional Animal Care and Use Committee (2015/SHS/1104).

### 2.2 Western diet and fructose model

Ten-week-old male C57Bl/6J mice were fed Western diet (D12079B; Research Diets), supplemented with 15% weight/volume fructose (Sigma-Aldrich, 57-48-7) in drinking water for 8 or 16 weeks whereas control mice received normal chow and tap water for 16 weeks. [14]

### 2.3 Lepr^db^ with MCD diet model

Male BKS.Cg-Dock7m+/+LeprdbJ (db/db) mice (Jackson Laboratory 009659) at 12 weeks of age were fed a normal chow diet or a MCD (A02082002BRi, Research Diets) diet for 2, 4 and 8 weeks to produce NASH stages. C57Bl/6J mice (NUSCARE C57BL/6 JInv) fed a normal chow diet served as control.

### 2.4 Liver-specific Dio1 Knockdown

Ten-week-old male C57Bl/6J mice were injected via tail vein with AAV8-ALB-eGFP-mDio1-shRNAmir or AAV8-ALB-eGFP-ctrl-shRNAmir (Lot 181231#13, Vector biolabs) and fed with NCD for two weeks followed by Western diet with fructose in the drinking water or NCD for next 12 weeks. A small group of mice (n=3) injected with the control shRNA were fed with NCD for reference purpose only and not used for statistical purposes.

### 2.5 Cell culture

AML12 cells were passaged in DMEM/F12 (cat. 11320-033), 10% FBS and 1x pen/strep, insulin transferrin selenium. 24 hours after plating the cells a mix of oleic acid 0.6M and palmitic acid (OAPA) in the above media with 1% BSA as carrier or only 1% BSA was added for 24, 48 and 36 hours. For siRNA transfection, AML12 cells were trypsinized, mixed with opti-MEM medium (Invitrogen, 31,985,070) containing Lipofectamine RNAimax (Invitrogen, 13,778,150) and Dio1 (ON-TARGET plus Mouse Dio1 (13370) siRNA SMARTPOOL (Dharmacon) or control siRNA (10 nM) according to the manufacturer’s recommendations. 24 hours later OAPA was added for 24 hours. The neutral lipid was stained with fluorescent dye BODIPY 493/503 (5 μg/ml) for 10 min. Oxygen consumption was measured at 37°C using an XF24 extracellular analyzer (Seahorse Bioscience Inc., North Billerica, MA, USA) [19].

### 2.6 Analysis

Triglyceride concentrations in the liver and serum (10010303; Cayman Chemical Company, Ann Arbor, MI) and total cholesterol (ab65390, abcam) were measured with colorimetric kits according to the manufacturer’s instructions after chloroform/methanol lipid extraction. Total RNA isolation was performed using InviTrap Spin Universal RNA kit (Stratec Biomedical), and RT-qPCR was performed as described previously [19] using QuantiTect SYBR Green PCR kit (Table primers in supplementary methods). Liver TH concentrations (T_4_ and bioactive T_3_) were measured by LC-MS/MS. Deiodinase 1 (Dio1) and 3 (Dio3) activity was measured by conversion of ^125^I-labelled rT_3_ and T_3_ respectively as previously described [12]. For western blot analysis, proteins were separated by SDS–PAGE under reducing conditions and transferred to nitrocellulose membranes. Membranes were blocked with 5% nonfat milk in phosphate-buffered saline with 0.1% Tween 20 (Sigma-Aldrich, P9416; PBST). The blots were incubated overnight at 4°C with primary antibodies. Immunoblot analysis was performed using an enhanced chemiluminescence procedure (GE Healthcare, RPN2106).

### 2.7 Statistical analysis

For the WDF model the groups were compared using a one-way ANOVA with a post-hoc Dunnett’s multiple comparison test to establish significance between the groups. For the *Lepr^db^* with MCD model the wildtype mice with a NCD diet were compared with the *Lepr^db^* mice on a NCD diet with an unpaired t-test to establish the effect of the genotype. To investigate the effect of the MCD diet the *Lepr^db^* on a NCD diet and 2,4 and 8 weeks of MCD were compared with a one-way ANOVA with a post-hoc Dunnett’s multiple comparison test to establish significance between the groups. Dio1LKD-WDF were compared to the control-WDF with an unpaired t-test. Data points smaller than Q1 – 1.5×IQR or greater than Q3 + 1.5×IQR were considered outliers and removed from further analysis. Prism 8 was used for statistical analysis. Data are represented in mean±SEM. Significance was established at p<0.05.

## 3. Results and Discussion

### 3.1 Dio1 expression and intrahepatic TH concentrations in mice fed Western diet and fructose (WDF)

To examine intrahepatic TH regulation during the progression of NAFLD, we employed two different NAFLD models to induce hepatosteatosis and NASH. In the first model, we fed mice a Western diet with 15% fructose water (WDF) diet for 8 and 16 weeks to induce steatosis and early-stage NASH and compared them with mice fed normal chow diet (NCD) (Fig. 1A) [14]. We observed that hepatic T_4_ decreased in mice fed WDF for 8 (15.3 pmol/g) and 16 (15.1 pmol/g) weeks compared to mice fed NCD (26.0 pmol/g) (Table 1). This decrease in hepatic T_4_ was not explained by decreased expression of the thyroid hormone transporters Mct8 and Mct10 as we found increased expression of both transporters in mice fed WDF for 8 weeks followed by a return at 16 weeks to levels similar to those in mice fed NCD (Table 1). In contrast to the prohormone T_4_, hepatic T_3_ was not significantly different in mice fed WDF for 8 weeks (2.1 pmol/g) and slightly decreased in mice fed WDF for 16 weeks (1.7 pmol/g) compared to mice fed with NCD (2.7 pmol/g) (Table 1). These data indicated intracellular regulation of T_3_ levels were likely mediated by deiodinases.

**Figure 1.**
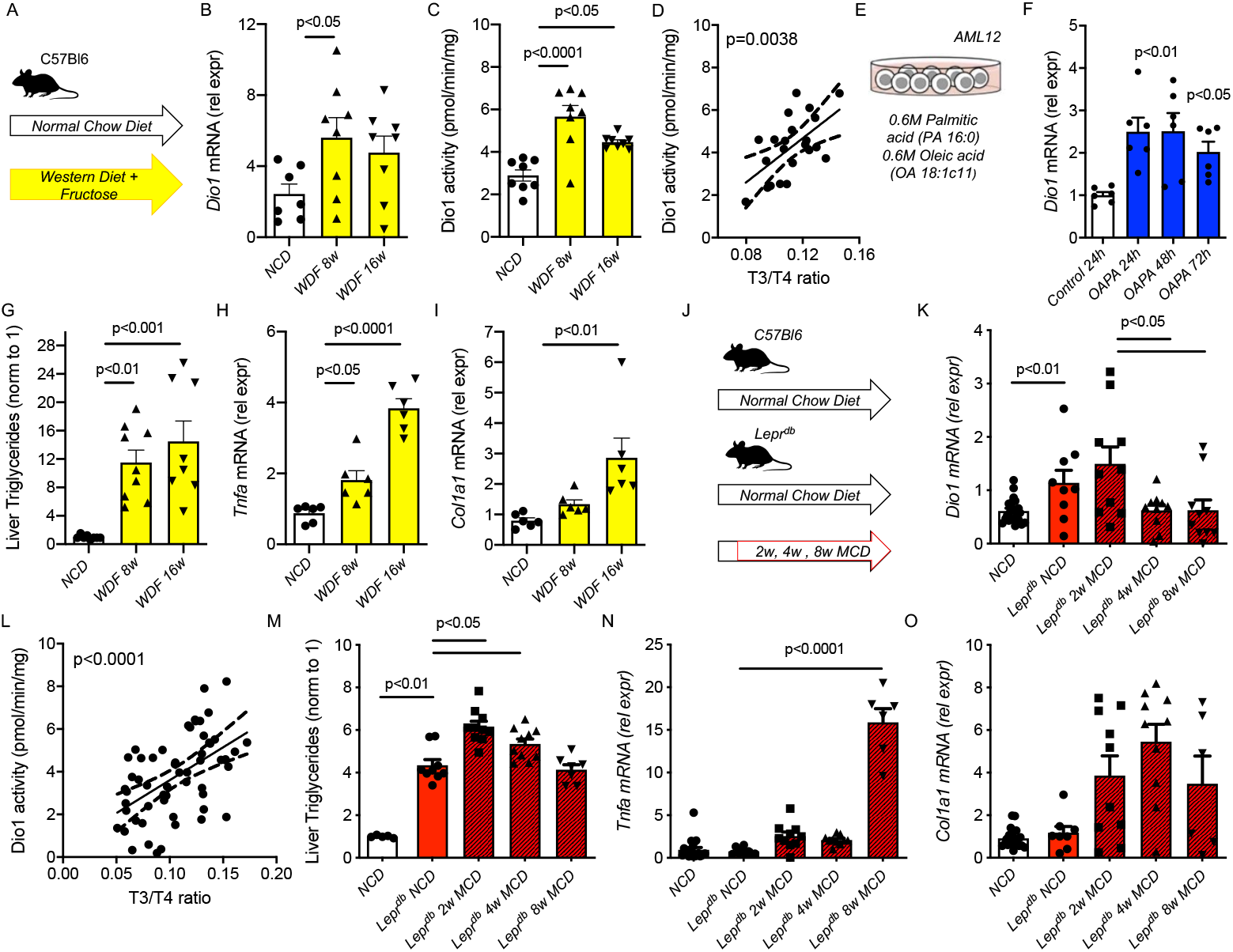
Dio1 increases early during the progression of NAFLD. (A) Western Diet with 15% fructose in the drinking water (WDF) for 8 and 16 weeks compared to Normal Chow Diet (NCD), (n=7-8 per group) (B,C) *Dio1* mRNA, DIO1 Activity in the WDF model, (D) Association between Deiodinase 1 activity and liver T_3_/T_4_ ratio in the WDF model (E) Mouse AML12 cell line with oleic acid and palmitic acid (OAPA) (6 wells per group) (F) *Dio1 mRNA* in the AML12 OAPA cell model (G-I) Liver triglycerides, *Tnfa mRNA* and *Col1a1 mRNA* in the WDF model (J) Lepr^db^ model with normal chow diet (NCD) or methionine and choline deficient diet (MCD) compared to C57Bl6 with NCD diet (n=6-10 per group) (K) *Dio1 mRNA* in the Lepr^db^ model (K) Association between Deiodinase 1 activity and liver T_3_/T_4_ ratio in the Lepr^db^ model (L-O) Liver triglycerides, *Tnfa mRNA* and *Col1a1 mRNA* in the Lepr^db^ model. Data is depicted in mean ± SEM.

**Table 1.** Liver parameters during the progression of NAFLD in the Western Diet with 15% Fructose in the drinking water (left) and the Lepr^db^ model with a methionine and choline deficient diet (MCD) or normal chow diet (NCD) (left). Significance of post-hoc analysis WDF 8w vs NCD and WDF 16 w vs NCD (left panel), *Lepr^db^* NCD vs C57Bl6 NCD, *Lepr^db^* 2w, 4w and 8w post-hoc analysis vs *Lepr^db^* NCD (right panel). * p<0.05, ** p<0.01, *** p<0.001, **** p<0.0001. Data is depicted in mean ± SEM.

|  | **WDF** |  |  | ***Lepr<sup>db</sup>* + MCD** |  |  |  |  |
| --- | --- | --- | --- | --- | --- | --- | --- | --- |
|  | NCD | WDF<br>8w | WDF<br>16w | Control<br>NCD | *Lepr<sup>db</sup>*<br>NCD | *Lepr<sup>db</sup>*<br>MCD<br>2w | *Lepr<sup>db</sup>*<br>MCD<br>4w | *Lepr<sup>db</sup>*<br>MCD<br>8w |
| Liver<br>T<sub>4</sub> (pmol/g) | 25.96<br>(2.20) | 15.29****<br>(1.02) | 15.08****<br>(0.82) | 29.11<br>(1.03) | 17.00**<br>(1.26) | 11.22**<br>(0.72) | 12.22***<br>(0.71) | 12.46***<br>(1.31) |
| Liver<br>T<sub>3</sub> (pmol/g) | 2.66<br>(0.23) | 2.09<br>(0.32) | 1.69*<br>(0.06) | 2.25<br>(0.18) | 1.99**<br>(0.09) | 1.48**<br>(0.09) | 1.3****<br>(0.09) | 1.55*<br>(0.16) |
| Dio3 mRNA<br>(rel expr) | 1.90<br>(0.42) | 2.55<br>(0.71) | 2.36<br>(0.39) | 2.09<br>(0.28) | 1.20<br>(0.29) | 2.93***<br>(0.33) | 1.64<br>(0.23) | 1.16<br>(0.27) |
| Dio3 activity<br>(fmol/min/mg) | 0.20<br>(0.03) | 0.14<br>(0.03) | 0.15<br>(0.01) | 0.13<br>(0.01) | 0.05****<br>(0.01) | 0.07<br>(0.01) | 0.11<br>(0.03) | 0.06<br>(0.02) |
| *Thrb* mRNA | 0.97<br>(0.14) | 0.94<br>(0.18) | 0.67<br>(0.10) | 3.21<br>(0.23) | 2.46<br>(0.27) | 2.66<br>(0.30) | 1.94<br>(0.21) | 3.86*<br>(0.57) |
| *Mct8* mRNA | 1.07<br>(0.27) | 2.72**<br>(0.43) | 1.94<br>(0.22) | 2.95<br>(0.32) | 2.35<br>(0.44) | 4.89*<br>(0.88) | 2.94<br>(0.42) | 6.82**<br>(1.34) |
| *Mct10* mRNA | 1.31<br>(0.36) | 3.75*<br>(1.01) | 1.20<br>(0.26) | 2.28<br>(0.38) | 1.16<br>(0.19) | 1.63<br>(0.34) | 1.31<br>(0.37) | 1.57<br>(0.33) |

We next examined Dio1 gene expression which is known to convert T_4_ to T_3_ in liver cells. *Dio1* mRNA increased more than 2-fold at both 8 and 16 weeks (Fig. 1B) in WDF mice. When Dio1 activity was measured by the conversion of ^125^I-labelled rT_3_, it increased significantly and was most pronounced at WDF 8 weeks compared to NCD mice (Fig. 1C). Increased Dio1 activity was associated with increased T_3_/T_4_ ratio for all mice indicating that it regulated intrahepatic T_3_ concentration (Fig. 1D). We also measured Dio3 mRNA and Dio3 activity, known to metabolize T_3_ to its inert metabolites, and found they were not significantly different in mice fed WDF *vs*. NCD (Table 1). To further examine *Dio1* mRNA induction during hepatosteatosis, we treated the mouse hepatic cell line, AML12, with 0.6M oleic acid and 0.6M palmitic acid (OAPA), and observed increases in *Dio1* mRNA expression at 24, 48, and 72 hours suggesting that this combination of saturated and monosaturated fatty acids could induce *Dio1* mRNA expression acutely in a cell autonomous manner (Fig. 1E, F).

We then measured hepatic triglyceride content and found that it was increased more than 10-fold in mice fed WDF for 8 and 16 weeks when compared to mice fed NCD (Fig. 1G). Hepatic tumor necrosis factor alpha (*Tnfa*) was slightly increased in mice fed WDF for 8 weeks and more than 3-fold in mice fed WDF for 16 weeks (Fig. 1H). Alpha-1 type I collagen (*Col1a1*) mRNA was significantly increased in mice fed WDF for 16 weeks (Fig. 1I). These data suggested that mice fed WDF for 8 weeks had hepatosteatosis and slight inflammation whereas mice fed WDF for 16 weeks developed early-stage NASH with induction of inflammation and fibrosis marker mRNAs. These changes occurred in parallel with increased *Dio1* mRNA expression and activity, and correlated with T_3_/T_4_ ratio. Taken together, these data suggested that although a previous report showed Dio1 decreased in late stage NASH [2], *Dio1* gene expression and activity increased in hepatosteatosis and early-stage NASH to maintain intrahepatic T_3_ concentration.

### 3.2 Dio1 expression and intrahepatic TH concentrations in Lepr^db^ mice fed a methionine and choline deficient diet

To validate this observation in a second NAFLD model, we used *Lepr^db^* mice that previously were shown to develop severe steatosis with only mild inflammation when fed normal chow diet (NCD) [15]. To induce the NASH phenotype, *Lepr^db^* were switched after 12 weeks of age to a methionine- and choline-deficient diet (MCD) for 2, 4, and 8 weeks (Fig. 1J) [15]. Lepr^db^ mice continued on NCD diet had hepatic T_4_ levels that were 42% lower than wild-type mice on NCD (Lepr^db^-NCD T_4_: 17 vs. control-NCD 29.11 pm/g). In this model, increases of the thyroid hormone transporter *Mct8* mRNA were found after 2 and 8 weeks in Lepr^db^ mice fed MCD diet. Next, we investigated intrahepatic T_3_ which decreased only by 12% in Lepr^db^-NCD compared to control-NCD (T3: 1.99 vs. 2.25 pmol/g). In this model we also observed an increase in Dio1 *mRNA* expression in steatotic livers (Lepr^db^-NCD *vs*. control-NCD) (Fig. 1K). Dio1 *mRNA* decreased below basal level in Lepr^db^ mice fed MCD for 4 and 8 weeks similar to NASH in rats previously observed by us [2]. The enzyme activity of Dio1 was positively correlated with the T_3_/T_4_ ratio (Fig. 1L) again showing a regulatory role of deiodinases in liver T3 availability.

Triglyceride content in livers from *Lepr*^db^-NCD mice increased more than 4-fold compared to control-NCD (Fig. 1M). Triglyceride content further increased in *Lepr^db^* fed MCD for 2 and 4 weeks. *Tnfa* and *Col1a1* mRNA were not significantly different in *Lepr*^db^-NCD and control-NCD mice. However, there was increased *Tnfa* mRNA expression starting at 2 weeks and continuing to 6 weeks suggesting ongoing inflammation after the initiation of MCD (Fig. 1N). *Col1a1* mRNA was increased more than two-fold at 2 and 4 weeks, and increased 16-fold at 6 weeks (Fig. 1). These findings suggested that fibrosis associated with gene expression became more prominent in *Lepr*^db^ mice fed MCD by 8 weeks. Taken together in this second NAFLD model also an early increase in Dio1 mRNA was observed associated with the T_3_/T_4_ ratio.

### 3.3 Liver-specific Dio1 shRNA knockdown in mice fed WDF

Since T3 stimulates fatty acid β-oxidation, we investigated whether Dio1 increases during hepatosteatosis and early-stage NASH were protective mechanisms to maintain intrahepatic T_3_ concentration in order to reduce triglyceride accumulation in the liver during nutritional overload. Accordingly, we prevented the early induction of Dio1 by liver-specific knockdown using shRNA against mouse Dio1 cloned under the control of mouse albumin promoter in an adeno-associated viral vector (AAV8-Albumin-eGFP-mDio1-shRNAmir). Two weeks after tail vein injection of shRNA, mice were started on WDF or NCD for 12 weeks before sacrifice (Fig. 2A). We observed that hepatic *Dio1* mRNA and Dio1 activity decreased by decrease 72% and 66%, respectively, in Dio1 KD mice fed WDF (Dio1LKD-WDF) compared to control mice fed WDF (control-WDF) (Fig. 2B). Control-WDF had increased body weight and fat mass compared to control mice fed NCD (control-NCD); however, there were no significant differences in body weight, fat mass measured by MRI, and food intake was similar for Dio1LKD-WDF and control-WDF mice (data not shown). Hepatic T_4_ levels were higher in Dio1LKD-WDF (40.6 pmol/g) than control-WDF as would be expected if Dio1 expression/activity were decreased (Table 2). Interestingly, hepatic T_3_ concentrations were not significantly different among the three groups of mice. When we analyzed sera from the mice, we saw increased T_4_ levels in Dio1LKD-WDF (72.4 nmol/l) compared to control-WDF (52.3 nmol/l). Previously, it was shown that whole body knockout of Dio1 increased serum T_4_ [16]. Our data showed that reduced hepatic Dio1 expression and activity was sufficient to exert this effect in mice. Of note, serum T_3_ levels were not significantly different among the three groups of mice. In this experiment we did not find any effect on expression of *Mct8* and *Mct10* mRNA levels (Table 2). Taken together, the Dio1LKD-WDF showed decreased *Dio1* gene expression and Dio1 activity resulting in reduced metabolism of intrahepatic T_4_ concentrations without any differences in body weight and food intake.

**Figure 2.**
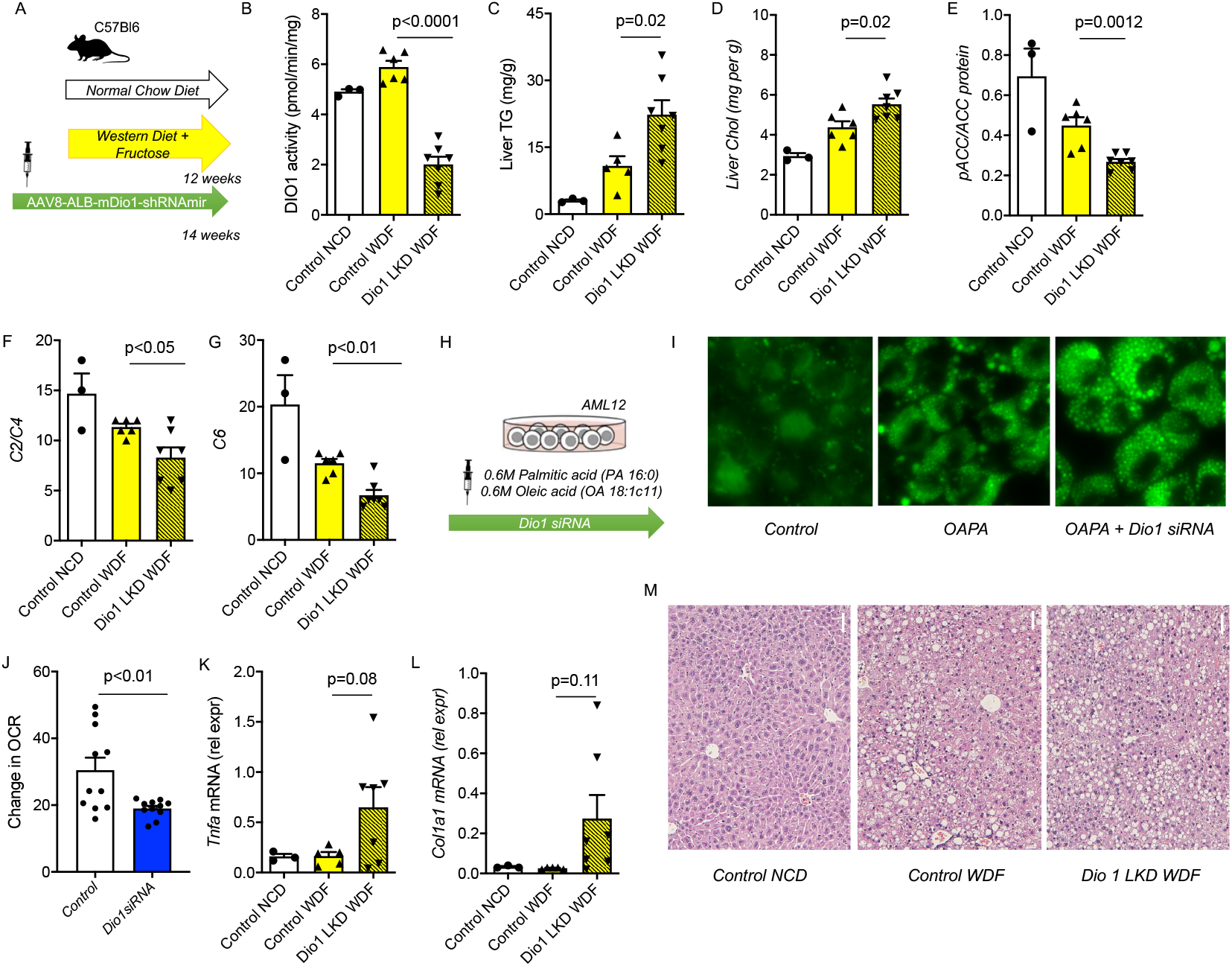
Dio1 KD increases liver triglycerides and cholesterol. (A) WDF model with injection of AAV8-Albumin-eGFP-mDio1-shRNAmir (WDF + Dio1 LKD) (n=7) or AAV8-Albumin-eGFP-ctrl-shRNAmir (WDF + control) (n=6). For reference a group of NCD + control shRNA was included (n=3). (B-G) DIO1 activity (B), liver triglycerides (TG; C), liver cholesterol (D), densitometric quantification of western blots analysing pACC/ACC (E), C2/C4 acylcarnitines (F), C6 acylcarnitines (G) in the WDF Dio1 KD model. (H) Schematic representation of in vitro experiment utilizing mouse AML12 cell line treated with oleic acid and palmitic acid (OAPA) combined with *Dio1* siRNA knockdown. (I) BODIPY staining of the AML12 cell line combined with OAPA and OAPA with Dio1 siRNA. (J) Change in oxygen consumption rate (OCR) after Dio1 siRNA in AML 12 cells. (K-L) *Tnfa mRNA* (K), *Col1a1* mRNA (L) in the WDF Dio1 KD model. (M) Histology of control NCD, control WDF, and Dio1 LKD WDF with TG content on average group. Data is depicted in mean ± SEM.

**Table 2.** Liver and serum parameters in normal chow diet (NCD) with a control shRNA (control-NCD) and western diet with 15% fructose in the drinking water (WDF) with a control shRNA (control-WDF) compared to Dio1 liver specific knockdown (Dio1LKD-WDF). Significance is shown for WDF control vs WDF Dio1 LKD. * p<0.05, ** p<0.01. Data is depicted in mean ± SEM.

|  | Control-NCD | Control-WDF | Dio1LKD-WDF |
| --- | --- | --- | --- |
| T<sub>4</sub> liver (pmol/g) | 41.40 (4.71) | 28.12 (2.40) | 40.59 (2.81)* |
| T<sub>3</sub> liver (pmol/g) | 6.07 (1.01) | 6.82 (1.06) | 8.86 (2.58) |
| T<sub>4</sub> serum (nmol/l) | 62.00 (4.04) | 52.33 (3.73) | 72.43 (3.09)** |
| T<sub>3</sub> serum (nmol/l) | 1.63 (0.05) | 1.68 (0.15) | 1.56 (0.10) |
| *Thrb* mRNA | 2.74 (0.55) | 1.13 (0.10) | 2.15 (0.31)* |
| *Mct8* mRNA | 3.67 (1.19) | 3.15 (1.41) | 4.36 (1.07) |
| *Mct10* mRNA | 1.23 (0.18) | 0.64 (0.09) | 1.20 (0.41) |

Dio1LKD-WDF showed increased hepatic triglyceride and cholesterol content compared with control-WDF (Fig 2 C,D). We next analyzed hepatic fatty acid metabolism and found Dio1LKD-WDF had a lower pACC/ACC protein ratio than control-WDF (Fig. 2E) suggesting there was increased fatty acid synthesis and decreased β-oxidation of fatty acids. We performed metabolomics of hepatic acylcarnitines as a measure of fatty acid β-oxidation. In control-WDF mice *vs*. control-NCD mice, there was a pattern of decreased short chain acylcarnitines (C2, C3, C4, C6) together with increased longer chain acylcarnitines (C10:1, C10:2, C12:1, C14:2) indicative of lower β-oxidation of fatty acids (Supplementary Figure 1). There was a further decrease in C2/C4 ratio and C6 in Dio1LKD-WDF compared to control control-WDF. We only observed an increase of the very long acylcarnitine C22:5 (Fig 2. F-G). We further examined the effects of Dio1 KD by siRNA *in vitro* combined with OAPA treatment in AML12 cells. We measured fat content by BODIPY staining in KD cells and observed increased fat content compared to control cells (Fig.2 H,I). Additionally, Dio1 KD cells exhibited decreased oxidative consumption rate by Seahorse analysis, consistent with lower fatty acid β-oxidation (Fig. 2J).

Last, we noticed there was a trend towards increased hepatic expression of TNFa and Col1a1 mRNA expression in Dio1LKD-WDF *vs*. control-WDF (Fig. 2K, L). When Dio1LKD-WDF were sacrificed 12 weeks after diet change, there was wide variability in the group at this transitional stage with some mice showing evidence of inflammation and fibrosis gene expression. In contrast, in this experiment none of the control-NCD or control-WDF had any increases in *Tnfa* and *Col1a1* mRNA at 12 weeks. Histology also showed increased fat droplets visible in Dio1LKD-WDF with increased ballooning of hepatocytes (Fig. 2M). Serum triglyceride, cholesterol and glucose levels were not significantly altered by Dio1LKD-WDF compared to control-WDF (data not shown).

## 4. Conclusions

Taken together, our study showed that blocking Dio1 induction increased hepatic triglyceride and cholesterol content, and caused a rapid progression towards NASH in several individual Dio1LKD-WDF mice. This finding may have important physiological ramifications and suggests that induction of Dio1 expression and activity in hepatosteatosis and early NASH may play a preventive role in NASH progression. Moreover, increased hepatic Dio1 in these early stages of NAFLD supports our previous observation that levothyroxine (T_4_) can be an effective treatment for hepatosteatosis since it can be converted to T_3_ intrahepatically [2]. Also, it is possible that patients with less induction of Dio1 in the early stages of NAFLD may be at higher risk for developing hepatosteatosis and NASH more rapidly. Decreased Dio1 is observed in older age, certain medications such as propranolol and propylthiohuracil, and selenium deficiency. Recently the first loss-of-function human Dio1 mutation causing changes in thyroid hormone metabolism was described [17]. On the other hand, a Dio1 polymorphism had increased Dio 1 activity [18]. Thus, epigenetic and genetic factors could alter Dio1 expression and/or activity, and thus affect the risk for progression of NAFLD. In conclusion, our findings show that hepatic Dio1 expression is sensitive to nutritional conditions and serves as a metabolic regulator during NAFLD progression to help decrease hepatosteatosis in early NASH.

## 5. Author contributions

**Eveline Bruinstroop:** Conceptualization, Methodology, Formal analysis, Investigation, Writing - Original Draft, Funding acquisition **Jin Zhou:** Conceptualization, Methodology, Investigation, Writing - Review & Editing **Madhulika Tripathi:** Investigation, Writing – Review & Editing **Winifred W. Yau:** Investigation, Writing – Review & Editing **Anita Boelen:** Conceptualization, Investigation, Writing – Review & Editing **Brijesh Kumar Singh:** Conceptualization, Methodology, Investigation, Supervision, Writing - Original Draft **Paul M. Yen:** Conceptualization, Methodology, Supervision, Writing - Original Draft, Funding acquisition.

## 6.

Acknowledgments

The authors would like to acknowledge Jia Pei, Keziah Tikno and An de Ruiter for their technical assistance and Eric Fliers for his critical appraisal of the manuscript. E.B. was funded by a Khoo Postdoctoral Fellowship Award, Niels Stensen Fellowship, Ter Meulen Grant of the Royal Netherlands Academy of Arts and Sciences, and Catherine van Tussenbroekfonds (A3-2). The authors would also like to acknowledge the funding from National Medical Research Council, Ministry of Health and A*STAR Singapore NMRC/OFYIRG/0002/2016 and MOH-000319 to B.K.S.; NMRC/OFYIRG/077/2018 to M.T.; and Clinician Scientist Award MOH-000306 to P.M.Y. The funding source had no involvement in study design; in the collection, analysis and interpretation of data; in the writing of the report; and in the decision to submit the article for publication.

