## Supplementary methods for "Early induction of hepatic deiodinase type 1 inhibits hepatosteatosis during NAFLD progression"

| | ***Forward*** | ***Reverse*** | ***Vendor*** |
|---|---|---|---|
| Thrb | GAGACTCTAACTTTGAATGGG | CGATCTGAAGACATTAGCAG | Sigma |
| Mct8 (Slc16a2) | CGTGCACCTGATGAAATATG | GATCATCATGGACATCAAGC | Sigma |
| Mct10 Slc16a10) | AAGCTCCATCGAGCCTCTGTA | GTCCCAAAATGACCAGTGACG | Sigma |
| Oatp1c1 | CCTTCTCTCTATCTGAGTCACGG | GGGCCATCCTTTACAGTCGG | Sigma |
| Dio1 | ACCCCCGATTCGCCCCTGACAA | ACCAGGGGCCTGCTGCCTTGA | Sigma |
| Dio3 | AAGAAAGTCAAAGGTTGTGG | AAAACGTACAAAAGGGAGTC | Sigma |
| Col1a1 | CGTATCACCAAACTCAGAAG | GAAGCAAAGTTTCCTCCAAG | Sigma |
| Actin | GTACCACCATGTACCCAGGC | AAGGGTGTAAAACGCAGCTC | Sigma |
